# Social task and aggressiveness shape immunity gene expression levels in the head of highly eusocial bees

**DOI:** 10.1101/2024.12.17.628725

**Authors:** Leonardo Campana, Carlos Antônio Mendes Cardoso Júnior, Anete Pedro Lourenço, Klaus Hartfelder

## Abstract

Living in crowded perennial nests, social insects encounter high risks of disease outbreaks and predation on colony resources. The innate immune system plays an important barrier against microbial pathogens, but its association with group defensive behaviours remains unexplored. Here we investigated the expression of innate immune system genes in the workers of three species of Brazilian stingless bees (*Scaptotrigona postica*, *Frieseomelitta varia*, and *Melipona quadrifasciata*), which differ in worker reproductive traits and colony defence strategies. First, we standardized a behavioural arena test to assess the species-specific individual aggressiveness level of the workers. Then, we quantified the transcript levels of four core genes of the Toll and Imd pathways in the heads and abdomens for the three main life cycle stages of the workers (nurse, guard and forager). For the heads we found a strong association between immunity gene expression and aggressiveness, with an upregulation in the heads of guards and foragers in comparison to their abdomen tissues of the aggressive species, *S. postica*. while the less aggressive species, especially *M. quadrifasciata* workers, did not show such strong upregulation. Our study establishes a potential molecular link between immunity pathways and complex social behaviours in the Brazilian stingless bees, which evolutionarily lost the sting apparatus for the defence of their colonies.

**BULLETED HIGHLIGHTS:**

- The expression of immune system genes increases with social task/age in the abdomen and head of highly eusocial bees.
- The expression of Toll and Imd pathway genes is upregulated in the heads of the more aggressive stingless bee species.
- In the more docile stingless bee species, immune gene expression is higher in the abdomen than in the head.
- Our findings suggest a possible involvement of immunity pathway signalling in the regulation of key social behaviours.

## INTRODUCTION

Social bees, including the honeybee and the stingless bees, are territorial animals that build perennial nests where they store food resources (honey and pollen) and rear and protect the brood and the highly reproductive queen. Such social life presents two types of risks. On the one hand, the nutritious resources in the colony can attract a variety of predators and nest parasites, and on the other, the large number of individuals in these nests represents an elevated risk for the spreading of diseases (Abbot, 2022). Yet strikingly, the analysis of the honeybee genome (The Honeybee Genome Sequencing Consortium, 2006) indicated a reduction in the number of genes comprising the bees’ immune response repertoire compared to other insects, such as *Drosophila melanogaster* and *Anopheles gambiae*. While this loss can be compensated by social behaviours (social immunity) that provide barriers to the spreading of diseases (Cremer et al., 2018), a reduction in the repertoire of immune system genes, however, was also observed in bee species with a solitary lifestyle (Barribeau et al., 2015), apparently representing an ancestral trait in the bees, and thus unrelated to the evolution of advanced sociality in this clade.

For the defence of their nests, social bees adopt different collective behaviours. In the honeybee, *Apis mellifera*, guard workers use the sting and also biting as the primary strategies against intruders (Abbot, 2022; Nouvian et al., 2016). In contrast, having evolutionarily lost their sting apparatus, the stingless bees (Meliponini) exhibit diverse defence strategies. Some launch mass attacks, biting invaders, some apply sticky resins on them, while others have nest entrances that are difficult to access (for reviews see Grüter, 2020; Roubik, 2023; Shanahan and Spivak, 2021). Also, guards can respond to invasion threats by attacking aggressively, releasing alarm pheromones, by forming a corridor of hovering guards, or by simply blocking a narrow nest entrance (Kärcher and Ratnieks, 2009).

Molecular signatures of the defensive (aggressive) behaviours against threats have been mainly studied in the Western honeybee, *A. mellifera*, and a strong correlation was found between the colony level of aggression and the frequency of certain alleles in the population (Avalos et al., 2020). At the individual level, workers were found to exhibit distinct gene expression profiles in their brains, differentiating more aggressive individuals from more docile ones (Rittschof, 2017). Such individual tendencies in aggressiveness are, however, masked by the prevailing behavioural disposition of the colony at any given moment, indicating that the colony genetic composition may override the individual workers’ genetic background in the decision to place or not an attack (Avalos et al., 2020). At the cellular level, a recent transcriptomics study (Traniello et al., 2023) revealed genomic subnetworks associated with colony aggression across various cell types and across brain regions involved in olfaction, vision, and multimodal sensory integration.

Previous studies have also demonstrated a correlation between aggressiveness and immune system activity. For instance, the highly aggressive Africanized hybrids of *A. mellifera* present strong traits of resistance against ectoparasites compared to more docile *A. mellifera* subspecies (Camazine, 1986). Also, relatively docile, healthy workers, as well as workers raised in less aggressive colonies, presented a gene transcriptional profile in the brain and in the abdominal fat body that was very similar to that of individuals suffering from acute pathogen infection (Rittschof et al., 2019), indicating that low aggression resembles a sickness-like behavioural state. Furthermore, transcriptional regulatory network analyses demonstrated that the neuro-genomic behavioural state of aggressive honeybee workers is characterized by high expression levels of the transcription factor Dorsal (Chandrasekaran et al., 2011).

Dorsal, a Nuclear Factor-kappa B (NF-kB) protein in insects, is a transcription factor that plays a key role in the Toll signalling pathway of the innate immune response of insects (Hoffmann, 1995), similar to NF-kB in the Toll-like receptors (TLR) signalling pathway in vertebrates (Imler and Hoffmann, 2001). Unlike vertebrates however, invertebrates rely solely on innate immunity as their defence against pathogens. This system consists of humoral and cellular responses mediated by haemocytes and antimicrobial peptides (AMPs) (Danihlík et al., 2015; Janeway and Medzhitov, 2002; Lourenço et al., 2018; Strand, 2008). In this context, the Toll signalling pathway plays a vital role in the immune response of insects. When activated in response to recognition proteins that detect specific components of the cell wall of fungi and Gram-positive bacteria, the Toll pathway triggers the translocation of the transcription factor Dorsal to the nucleus, leading to the transcription of AMP genes (Evans et al., 2006; Kingsolver et al., 2013; Zhang et al., 2021).

Similarly, the Imd signalling pathway is triggered by the recognition of peptidoglycans from Gram-negative bacteria, resulting in the activation of the NF-kB transcription factor Relish and, consequently, the expression of specific AMPs (Zhai et al., 2018). In bees, the Toll and Imd pathways are known to regulate the expression of AMPs, such as Abaecin and Defensin-1 (Lourenço et al., 2013, 2018). These pathways are, however, not restricted to immune functions. In fact, as indicated by the primary name ‘Dorsal’, this gene was first identified in *Drosophila* embryonic mutant screens as a key transcription factor in the determination of the dorsal-ventral embryonic axis (Lynch and Roth, 2011).

Both, innate immunity and aggressive behaviours are aspects related to somatic maintenance, and both are energetically costly and, thus, likely of relevance to trade-offs in life histories (Flatt et al., 2013; Stearns, 1992). In general, reproductive activity and immune responses frequently show a negative correlation in insects, a pattern observed in both solitary species from orders such as Orthoptera, Diptera, Odonata, Hemiptera, and Coleoptera, as well as in some social insects, such as certain ant species (Schwenke et al., 2016). But how does this correlation look like in social insects, especially in the highly eusocial ones, where colony (somatic) maintenance and reproductive activity are dissociated into two morphologically distinct castes, the workers and the queen?

Studies on social insect immunity, colony defence strategy, reproductive activity, and their associated gene networks have been mainly conducted using *A. mellifera* workers as a model. However, the honeybees (Apini) represent a single genus (*Apis*) consisting of less than a dozen species, and thus are a relatively small and rather homogeneous group in the phylogeny of the bees (Apidae). In contrast, the stingless bees (Meliponini) comprise over 500 recognized species distributed among 58 genera (Grüter, 2020; Melo, 2021). With a pantropical distribution, they exhibit a high level of biological diversity in terms of nesting sites, reproductive strategies, and nest defence. Stingless bees, like honeybees, are threatened by anthropogenic impacts, such as crop monoculture and pesticide use. However, unlike honeybees, they thrive in greenhouse environments, where they are increasingly recognized as efficient pollinators in high-value agriculture. (Grüter, 2020).

Taking advantage of the stingless bees’ biological variability, we set out in this study to investigate the relationship between aggressiveness (defence behaviour) and the immune system profile of the workers of three stingless bee species with distinct life history configurations (Fig. 1). The workers of *Scaptotrigona postica* are very aggressive in their nest defence and are able to lay eggs in the presence of the queen (Nogueira-Neto, 1997). The workers of *Melipona quadrifasciata* are also able to lay eggs in the presence of the queen, but they are notably docile (Grüter, 2020; Nogueira-Neto, 1997), and the workers of *Frieseomelitta varia,* while moderately defensive, are completely sterile (Boleli et al., 1999; Nogueira-Neto, 1997). Hence, we considered that this contrast in character combinations would allow us to address the question as to how life histories and their respective trade-offs (Stearns, 1988), specifically those between the workers’ aggressiveness in the nest defence and their reproductive potential became configured with respect to a costly system, such as the innate immune response. Specifically, while on the one hand the investment in aggressiveness is likely very costly to the individual worker, but benefits the colony fitness, worker reproduction on the other hand creates a fitness conflict, primarily with the queen, but in the honey bee, also with the other workers (Ratnieks et al., 2006).

**Figure 1.**
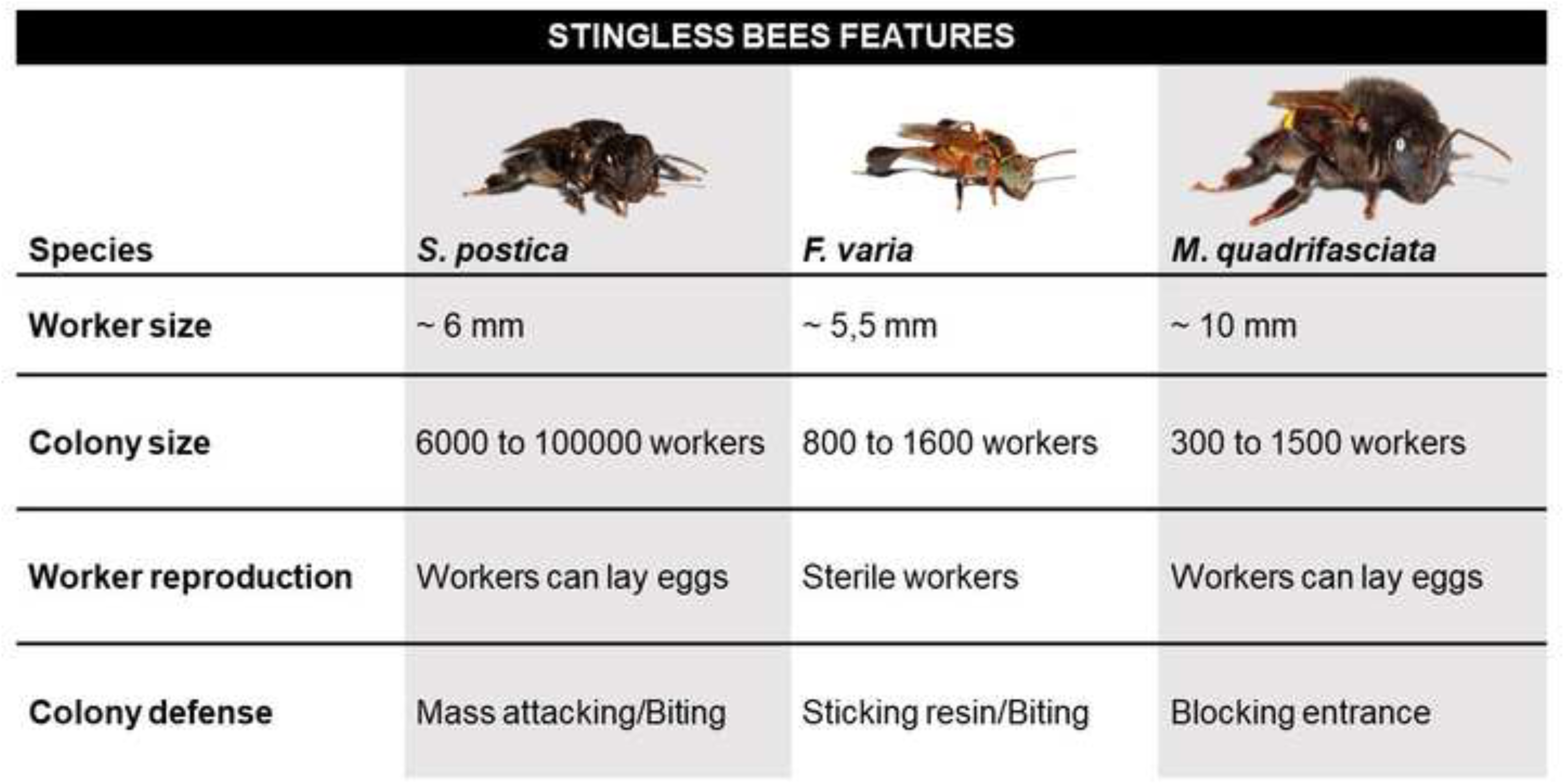
The stingless bee species investigated in this study and their major life history traits. Images are credited to Dr. Cristiano Menezes.

Hence, the underlying hypotheses that we proposed to address were that: (a) even in the absence of a specific immune challenge, the expression levels of immunity-related genes increase with the age of the bees, reflecting their progression in social behaviours (Wilson-Rich et al., 2008), and (b) the expression levels reflect the level of aggressiveness and the reproductive biology of the workers. To do so, we adapted an arena assay that allowed us to quantitatively assess the species-specific levels of aggressiveness for individual workers, and we evaluated the expression levels of a set of four key genes of the Toll and Imd pathways in the heads and abdomens for three important behavioural stages in the workers’ life cycle. Finally, we compared the results obtained for the three stingless bees with those of a corresponding analysis that we performed on honeybee workers.

## MATERIAL AND METHODS

### The arena behavioural assay: A quantitative tool for assessing the species-specific aggressiveness level of individual workers

To obtain information about the aggressiveness at the individual level for the workers of the three focal stingless bee species, we performed an arena assay based on the protocol of Dierick (2007) for *Drosophila melanogaster*, which has subsequently been applied to honeybees by Shpigler et al. (2017). This assay eliminates variables associated with the social colony context, including the presence of alarm pheromones elicited by nestmates, as well as environmental factors, such as ambient temperature, group size, diet, time, and territorial size in the bees’ reaction against a perceived threat.

Workers of *Scaptotrigona postica* (Latreille, 1807), *Frieseomelitta varia* (Lepeletier, 1836), and *Melipona quadrifasciata* (Lepeletier, 1836) were collected from one single colony of each species, all maintained in the apiary at the Department of Genetics, Ribeirão Preto Campus, University of São Paulo, Brazil. From each colony of *S. postica* and *F. varia* (Lepeletier, 1836), 100 newly-emerged workers and 10 nurse-stage workers were collected and kept together in a single cage (8 x 11 x 13 cm) in an incubator at 34 °C for 14 days. For *M. quadrifasciata*, due to their larger body size, the number of newly emerged workers (N = 40) and nurse-stage workers (N = 4) in the cage was reduced. The cages were provided daily with pollen collected from the colonies of origin and with commercial *A. mellifera* honey diluted in water at 50%. The presence of nurse bees in the cages is important, as they feed the newly emerged workers during their first days of adult life.

After 14 days in the cages, the workers were submitted to a 15-hour food restriction period prior to the arena assays. This food restriction period proved to be essential, as, during a pilot experiment, we had noted, in the case of all three species, that workers that were fed until the day of the assay displayed a disregard for the presence of an intruder in the assay arena (Fig. S1). The 15-days-old workers were then individually placed into 9 cm diameter Petri dishes lined with filter paper. After one minute of acclimatization, an intruder, a newly emerged *Tetragonisca angustula* worker, was introduced into each dish. The workers of the stingless bee species *T. angustula* are smaller than those of our focal species, and not aggressive when they are still young. Each Petri dish was filmed for 5 minutes, and the recordings were then analysed using Boris v. 8.11.3 software (Friard and Gamba, 2016). In total, we performed arena assays for 42 *S. postica* workers, for 42 *F. varia* workers, and 19 *M. quadrifasciata* workers.

The following aggression parameters were considered: (i) Antennation (events of touching the intruder with the antennae), (ii) frequency of attacking (percentage of tested individuals that show any attack behaviour against the intruder during the observation period), (iii) latency to attack (time until the first attacking event occurred), (iv) the attacking index (total amount of time spent attacking the intruder during the five-minute period), and (v) the pursuit index (total amount of time spent pursuing the intruder during the five minutes).

### Sampling of the worker bee life cycle stages for gene expression analysis

Workers of *S. postica*, *F. varia*, and *M. quadrifasciata* were obtained from the same colonies as those used for the arena aggression assays. For each species, sampling was done from three colonies. From each colony, seven workers were retrieved for each of the three life cycle stages: nurse, guard, and forager. Sampling was done during the Southern summer season, between November 2021 and March 2022, always between 9:00 and 10:00 am. Workers of the corresponding life cycle stages were also sampled from a honeybee colony kept in the same apiary (*Apis mellifera*, Africanized hybrids) in August 2022.

The sampling procedure was as follows: First, guards were collected, considering that these were the bees that left the nest and attacked the collector’s entomological sweep net after three knocks with a hive chisel on the front board of the hive. Guards of *S. postica* and *M. quadrifasciata* generally had an empty pollen basket (corbicula). In contrast, the workers of *F. varia* that attacked the net all had their corbiculae (pollen baskets on the hind legs) full of resin, as part of their colony defence strategy (Roubik, 2023). Next, foragers were collected in front of the hives, when returning from the field with pollen-filled corbiculae. Finally, the colonies were opened, and nurse-stage bees were collected from the brood area. In all three species, nurse-stage bees can be identified by their enlarged abdomen, due to their larval food-filled crop, and, their lighter body colour, as they are still younger than the guards or foragers, and not yet fully pigmented (Nogueira-Neto, 1997). Nurse bees are primarily engaged in the provisioning of newly constructed brood cells with larval food. Fig. S2 shows the phenotypic characteristics of these life cycle stages for each of the three species.

After sampling, the bees were briefly put on ice to diminish their motility and then snap frozen in liquid nitrogen. The heads and abdomens were separately transferred into Eppendorf vials containing 300 µL of TRIzol reagent (Invitrogen, Waltham, MA, USA) and stored at −80 °C until RNA extraction. For the two species of smaller body size, *S. postica* and *F. varia*, each sample was composed of a pool of heads or abdomens from three individuals, while for the larger species *M. quadrifasciata* and *A. mellifera*, each sample consisted of a single individual only. Pooling was necessary to obtain the RNA yield required for gene expression analysis.

In this respect, we recognize as a limit of our study that we not separate between age and social function. Separating these would require the setup of single-cohort colonies, but such an experimental design is currently only available for honey bees (Marco Antonio et al., 2008; Sullivan et al., 2000) and not for stingless bees. While in the current study our emphasis was primarily on social task and not on age, the compiled bionomics data available for stingless bees (Grüter, 2020) show that there is a clear age sequence in social task performance.

### RNA extraction and gene expression analysis

The head and abdominal tissues were macerated and homogenized with a pestle while still in the TRIzol reagent before proceeding with the RNA extraction according to the manufacturer’s protocol. RNA extracts were treated with RNase-free DNase (Invitrogen) to remove residual DNA. The quantity and quality of the RNA was checked spectrophotometrically (NanoVue, GE Healthcare, Chicago, IL, USA). Reverse transcription was performed using 1 μg of total RNA for cDNA synthesis, using the Ultrascript 2.0 cDNA synthesis kit (PCRBiosystems, London, UK), following the manufacturer’s instructions.

The transcript levels of *Toll*, *Dorsal*, *Myd88* (Toll signalling pathway), and *Imd* (Imd signalling pathway) were assessed through quantitative real-time PCR (RT-qPCR) assays. Primer sequences for these genes were based on the respective genomic sequences of *M. quadrifasciata* and *F. varia* (Table S1). The same primers were also tested and successfully used to amplify the corresponding transcripts of *S. postica,* for which a sequenced genome is not yet available.

Primer efficiencies were calculated as E = 10^[-1/slope]^, using a 1∶10 dilution series of a cDNA pool containing equal amounts of RNA from all experimental groups and species (Table S1). This pooling approach is a widely accepted method to estimate general primer performance across conditions. The primers for the endogenous control genes *Rpl32* and *Rps18* had previously been designed and validated for *F. varia and M. quadrifasciata* (Freitas et al. (2019). Additionally, as this was the first time these reference genes were used here also for *S. postica*, we confirmed their suitability by a series of additional efficiency tests using individual cDNA pools from each experimental group (head and abdomen tissues from both guards and foragers). The results show their high and consistent efficiencies (Table S1). For the normalization process, the Ct results of the two reference genes were combined by calculating their mean, and this average Ct value was used. The primer sequences for the corresponding *A. mellifera* immune system genes are based on the work by Evans et al. (2006), and those for the honeybee endogenous control gene *Rpl32* (*Rp49*) on Lourenço et al. (2008).

Amplification mixtures were set up in a total volume of 10 μL, containing 5 µl of the 2×qPCR SyGreen Mix kit (PCR Biosystems, London, UK), 1 µL of 10 × diluted cDNA, primers (forward and reverse) at a final concentration of 0.125 pmol, and ultrapure water completing the final volume. The assays were performed in a StepOnePlus real-time PCR system (Applied Biosystems, Waltham, MA, USA) with the following amplification protocol: 95 °C for 10 min, followed by 40 cycles at 95 °C for 15 s and 60 °C for 30 s. After these cycles, a melting curve analysis was performed to confirm the specificity of the amplification products. Assays were done in biological septuplicates (n = 7 per colony), and each sample was analysed in technical triplicates. The relative expression of the genes of interest was calculated by the 2^-ΔΔCt^ formula proposed by Livak and Schmittgen (2001).

To ensure the robustness of our analyses, we decided to remove the nurse bee samples of *S. postica* from the analyses, as for this group the reference genes exhibited considerable variation compared to the guards and foragers (Fig. S3). Despite this adjustment, it is still possible to compare gene expression profiles between species with reproductive (*M. quadrifasciata*) and non-reproductive (*F. varia*) workers, while the *S. postica* data were used to specifically discuss the relationship between the life history traits aggression and immunity.

### Statistical analysis

Statistical analyses were performed in the RStudio platform (R version 4.3.0). Data obtained from the arena assays were submitted to Shapiro-Wilk and Kolmogorov-Smirnov tests to check the adequacy for parametric testing. Levene’s test was used to verify homoscedasticity. As these conditions were not met, we used the Kruskal-Wallis test (dplyr package) with Dunn’s post-hoc test for multiple comparisons of three or more groups. For the analysis of the gene expression data, we used a Generalized Linear Mixed Model (GLMM) (lme4 package), considering “body compartment” and “life cycle stage” as fixed factors and “colony” as random. For *Apis mellifera*, in the absence of colony replicates, we utilized the GLM test (lme4 package) to identify differences between means. The log link function was used to approximate the gene expression data to a Gaussian distribution, which was checked by inspecting the residuals (DHARMa package). Graphical representations of the data were generated with GraphPad Prism v. 7 (GraphPad Softwar, Boston, MA). An adjusted *p*-value < 0.05 was considered significant in all analyses.

### Ethics statement

For the behavioural assay, the bees were kept in social groups in cages of suitable size and with *ad libitum* access to food. The 15-hours food restriction period had no visible negative effect on the well-being of the bees. After the arena assay, the bees were sacrificed by instant freezing in liquid nitrogen, as they could not be returned to their colonies without being attacked there as intruders. For the gene expression analyses the bees were briefly cooled on ice to diminish their motility before instantaneous freezing in liquid nitrogen. This is a standard procedure to calm down before introducing them into any experimental procedure.

The maintenance of the native stingless bee colonies in the apiary of the Ribeirão Preto Campus of the University of São Paulo is authorized by the *Instituto Chico Mendes de Conservação de Biodiversidade – Ministério de Meio Ambiente*, Brazil, under the licence 81843-1 to the senior author.

## RESULTS

### Species-specific individual aggressiveness of stingless bee workers

The arena assays demonstrated that under standardised conditions (number of bees of similar age, temperature, and diet) *S. postica* showed the highest scores in the aggression indices, while *M. quadrifasciata* exhibited the lowest scores (Fig. 2, Table S2). The scores for *F. varia* indicated intermediate levels of aggressiveness. *S. postica* was the species with the highest frequency of individuals that attacked the intruder in the arena. This species also was the one with the highest frequency of fatal attacks (Fig. 2A). In comparison, the workers of *F. varia* and *M. quadrifasciata* had much lower attacking frequencies. The behavioural profiles of the individual bees inside the arena were also different for the three species. Already within the first minute after introducing the intruder specimen, the workers of *S. postica* behaved agitated, walking around quickly, while those of *F. varia* took short flights, during which they collided with the intruder or the arena wall. *M. quadrifasciata* workers, in turn, appeared rather lethargic, with little movement in the arena.

**Figure 2.**
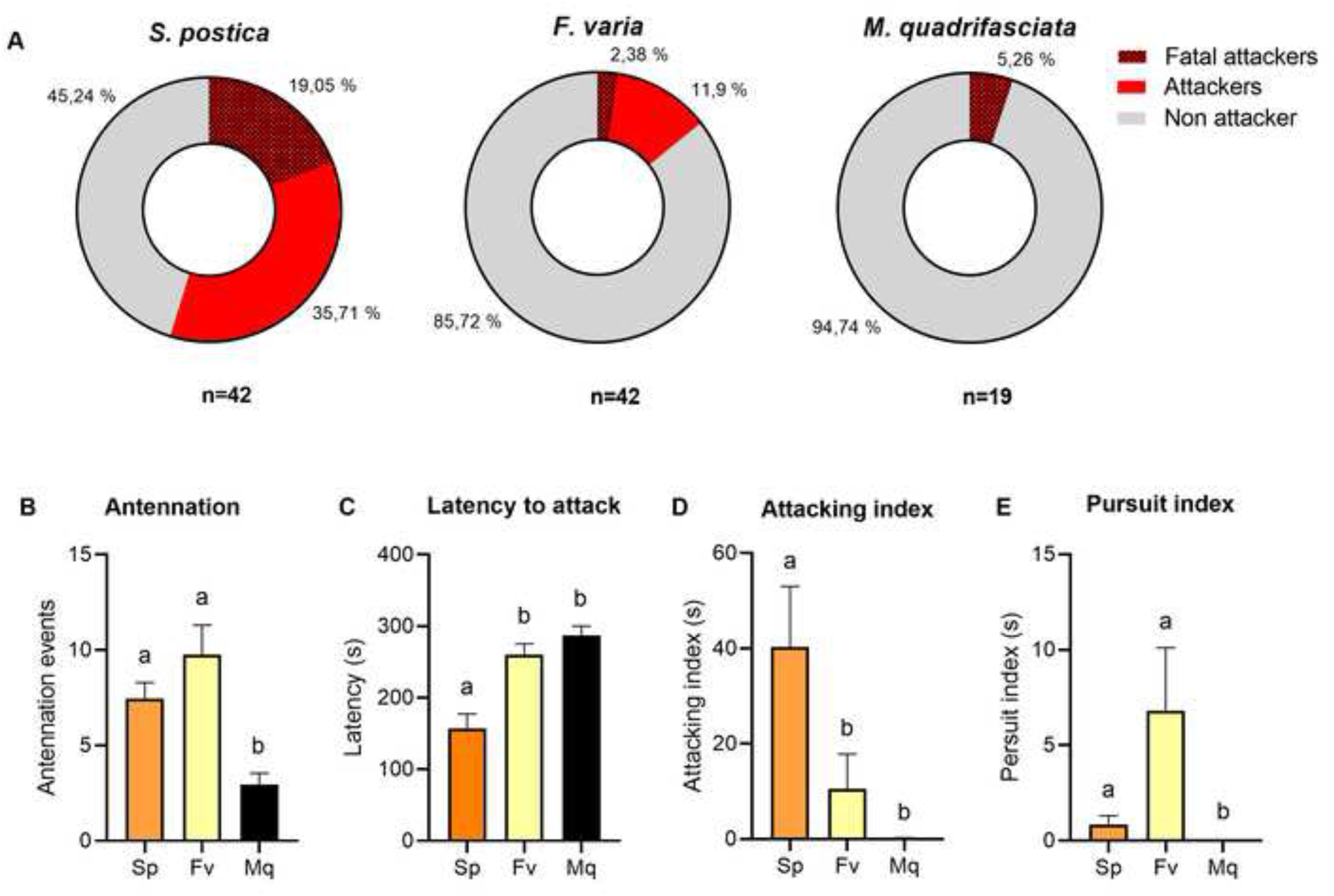
Arena test results on aggressiveness of individual workers of three stingless bee species. (A) Attack frequency and fatal attack frequencies for *S. postica*, *F. varia*, and *M. quadrifasciata*. The dark red colour represents the percentage of fatal attackers, i.e., those that killed the intruder, the red colour represents the percentage of attackers that did not kill the intruder, and the grey colour represents the percentage of bees that did not attack the intruder. (B) Number of antennation events: *S. postica* and *F. varia* exhibit have higher number of antennation events when compared to *M. quadrifasciata* (Kruskal-Wallis: *H =* 20.02, *P <* 0.001). (C) Latency to attack: *S. postica* exhibits the shortest latency to attack compared to *F. varia* and *M. quadrifasciata,* with no significant differences among the latter two species (Kruskal-Wallis: *H =* 21.05, *P* < 0.001). (D) Attacking index: *S. postica* has a higher attacking index than *F. varia* and *M. quadrifasciata* (Kruskal-Wallis: *H =* 23.31, *P* <0.001). (E) Pursuit index: All three species showed differences in the chasing index (Kruskal-Wallis: *H* = 6.09, *P* < 0.05); however, pairwise comparisons revealed no significant differences. Orange: *S. postica* (Sp), Yellow: *F. varia* (Fv), Black: *M. quadrifasciata* (Mq). Different letters indicate statistically significant differences among the three species. The error bars represent the standard error of the mean (SEM).

All workers of *S. postica* and *F. varia*, even the non-attacking individuals, exhibited curiosity towards the intruder, with several events of observed antennation. In contrast, the workers of *M. quadrifasciata* did not engage with the intruder, although they were active and moving around the arena like the other species. (Fig. 2B). The only attack event that we saw for an *M. quadrifasciata* worker was by an individual that behaved very agitated already from the start of the observation period, heavily beating its wings and vibrating its body.

The species with the shortest attack latency towards the intruder, that is, it attacked the intruder faster than the other species, was *S. postica* (Fig. 2C: Kruskall-Wallis test: *H =* 21.5, *P* < 0.01). *F. varia* and *M. quadrifasciata* showed longer attack latencies. The attacking index, which is the time the individual remained attacking the intruder, was higher for *S. postica* than for *F. varia* and *M. quadrifasciata* (Fig. 2D: Kruskal-Wallis test: *H* = 23.31, *P* < 0.001), and among the three species, *S. postica* and *F. varia* workers presented the highest number of pursuit events (Fig. 2E: Kruskal-Wallis test: H = 6.094, P < 0.5). During these events, *F. varia* workers often pursued the intruder with opened wings, a behavior not observed in *S. postica* workers. However, these short flights were not counted as pursuit events; only pursuits that occurred while walking on the plate were considered, to maintain the same standard for all three species. No pursuit event was observed in the case of *M. quadrifasciata*. In these events, the *F. varia* workers used to pursue the intruder with opened wings, a behaviour that was not seen in *S. postica* workers. No pursuit event was observed in the case of *M. quadrifasciata*.

As the arena test permitted us to quantify the individual aggression levels of the three species under a controlled experimental setup, our results provide evidence that *S. postica* workers exhibit higher levels of aggression compared to the other species. *M. quadrifasciata* workers are rather docile, and *F. varia* workers show an intermediate level of aggressiveness. While this confirms the common impression one typically has when working with colonies of these species, the arena test results can now provide quantitative indices for the individuals’ species-specific aggressiveness levels, independent of eventually variable colony conditions, which cannot be easily standardized in the field. Of note, we did not include *A. mellifera* workers in the arena test because their defensive behaviour differs significantly from that of stingless bees. While stingless bees primarily rely on biting as a means of defence, *A. mellifera* workers typically attempt to sting, rarely engaging in biting. This fundamental difference in behaviour would require distinct criteria for quantifying aggression, making direct comparisons between the species problematic. Additionally, *A. mellifera* workers do not survive sufficiently long in confinement due to their inability to defecate without flying, leading to physical debilitation and mortality over time. In contrast, stingless bees can defecate within the cage and even organize waste in specific areas, allowing for easier maintenance and ensuring their survival in standardized experimental setups (See Discussion for details).

### Body compartment and social task-specific levels of immunity gene expression in the workers of stingless bees

The three stingless bee species presented distinct profiles of immunity gene expression levels for the head and abdomen. For *S. postica*, the most aggressive of the three species, we found that the workers showed elevated transcript levels of *Toll* (GLMM: χ^2^ = 109.6, *P* < 0.001), *Myd88* (GLMM: χ^2^ = 25.9, *P* < 0.001), *Dorsal* (GLMM: χ^2^ = 34.8, *P* < 0.001), and *Imd* (GLMM: χ^2^ = 30.1, *P* < 0.001) in the head when compared to the abdomen (Fig. 3, Table S3). For the *Myd88* gene, the guards’ heads actually showed higher expression when compared with the foragers’ heads (GLMM: *t* = −3.5, *P*= <0.001). For *M. quadrifasciata*, the most docile of the three species, the directionality was exactly the opposite, with significantly higher levels for *Toll* (GLMM: χ^2^ = 133.1, *P* < 0.001), *Myd88* (GLMM: χ^2^ = 4.2, *P* = 0.03), and *Dorsal* (GLMM: χ^2^ = 11.5, *P* < 0.001) in the abdomen (Fig. 4, Table S4). *F. varia* was similar to *S. postica* regarding the expression of the genes *Myd88* (GLMM: χ^2^ = 41.6, *P* < 0.001) and *Imd* for guards and foragers (Fig. 5, Table S5: GLMM: χ^2^ = 19.7, *P* < 0.001), but the difference between the two body compartments was much less pronounced.

**Figure 3.**
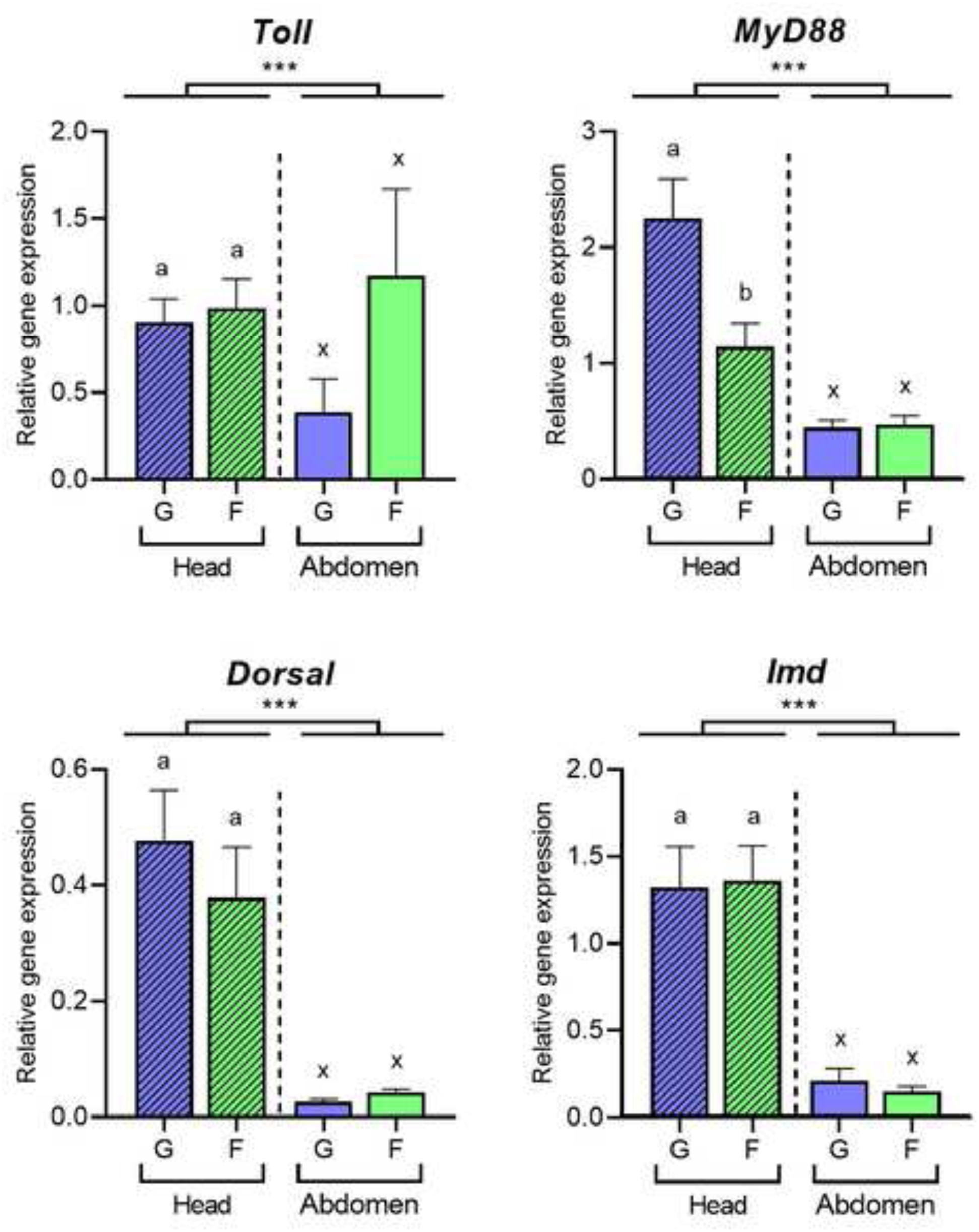
Relative expression levels of four immune system genes in the heads and abdomens of *Scaptotrigona postica* guards and foragers. Bar plots representing the transcript levels for (A) *Toll,* (B) *Myd88*, (C) *Dorsal,* and (D) *Imd*. Blue: Guard (G) and Green: Forager (F). Different letters indicate statistically significantly differences among the three live cycle stages within each body compartment. Asterisks indicate statistical differences between the two body compartments (GLMM). Symbols (***) *P* < 0.0001, (**) *P* < 0.001, (*) *P* < 0.05, (n.s.) non-significant. Sample size per colony, *N* = 7; per life cycle stage within each body compartment, *N* = 21 (3 colonies). The error bars represent the standard error of the mean (SEM).

**Figure 4.**
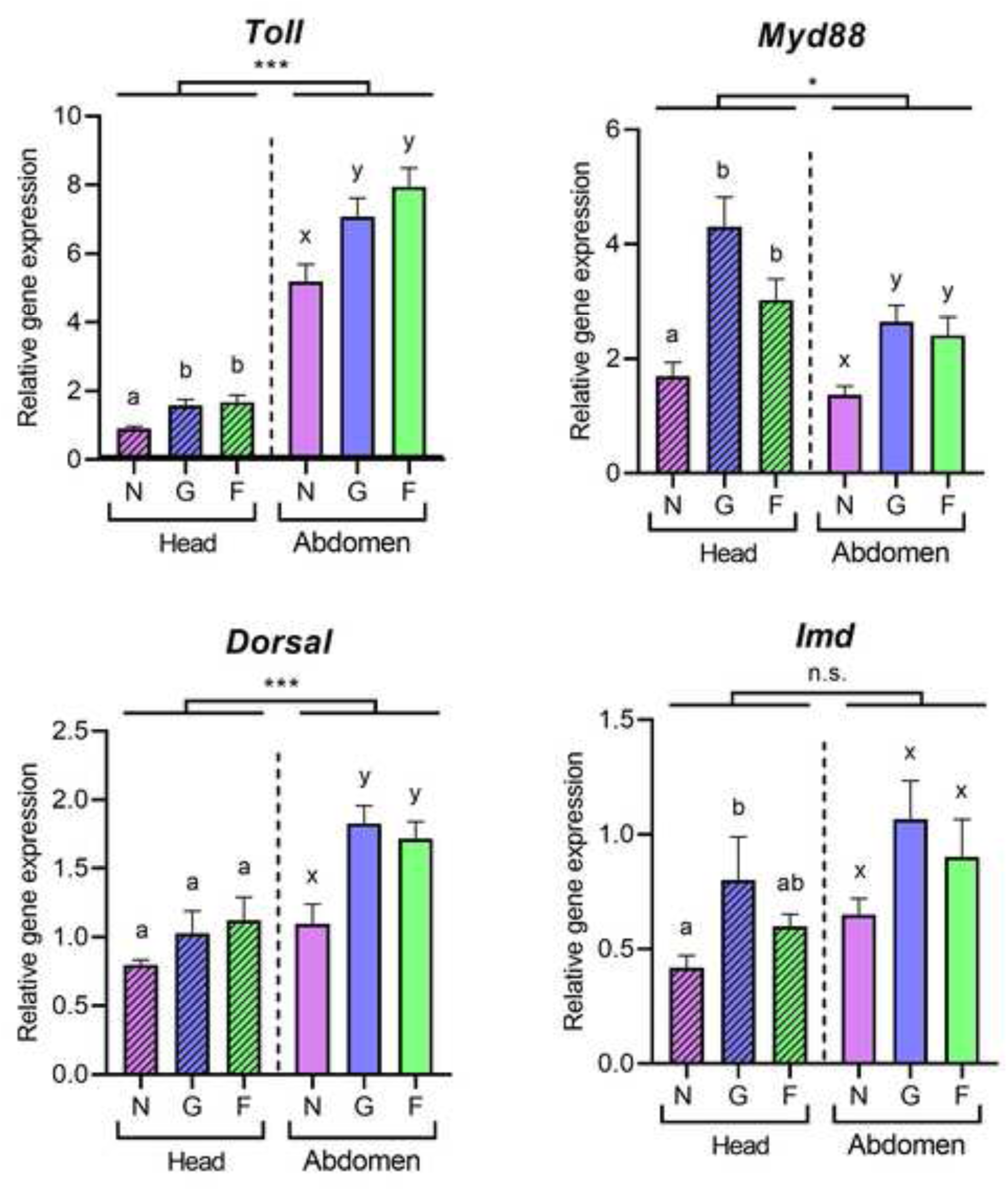
Relative expression levels of four immune system genes in the heads and abdomens of *Melipona quadrifasciata* nurses, guards, and foragers. Bar plots representing the transcript levels for (A) *Toll,* (B) *Myd88*, (C) *Dorsal,* and (D) *Imd*. Purple: Nurse (N), Blue: Guard (G), Green: Forager (F). Different letters indicate statistically significantly differences among the three live cycle stages within each body compartment. Asterisks indicate statistical differences between the two body compartments (GLMM). Symbols (***) *P* < 0.0001, (**) *P* < 0.001, (*) *P* < 0.05, (n.s.) non-significant. Sample size per colony, *N* = 7; per life stage within each body compartment, *N* = 21 (3 colonies). The error bars represent the standard error of the mean (SEM).

**Figure 5.**
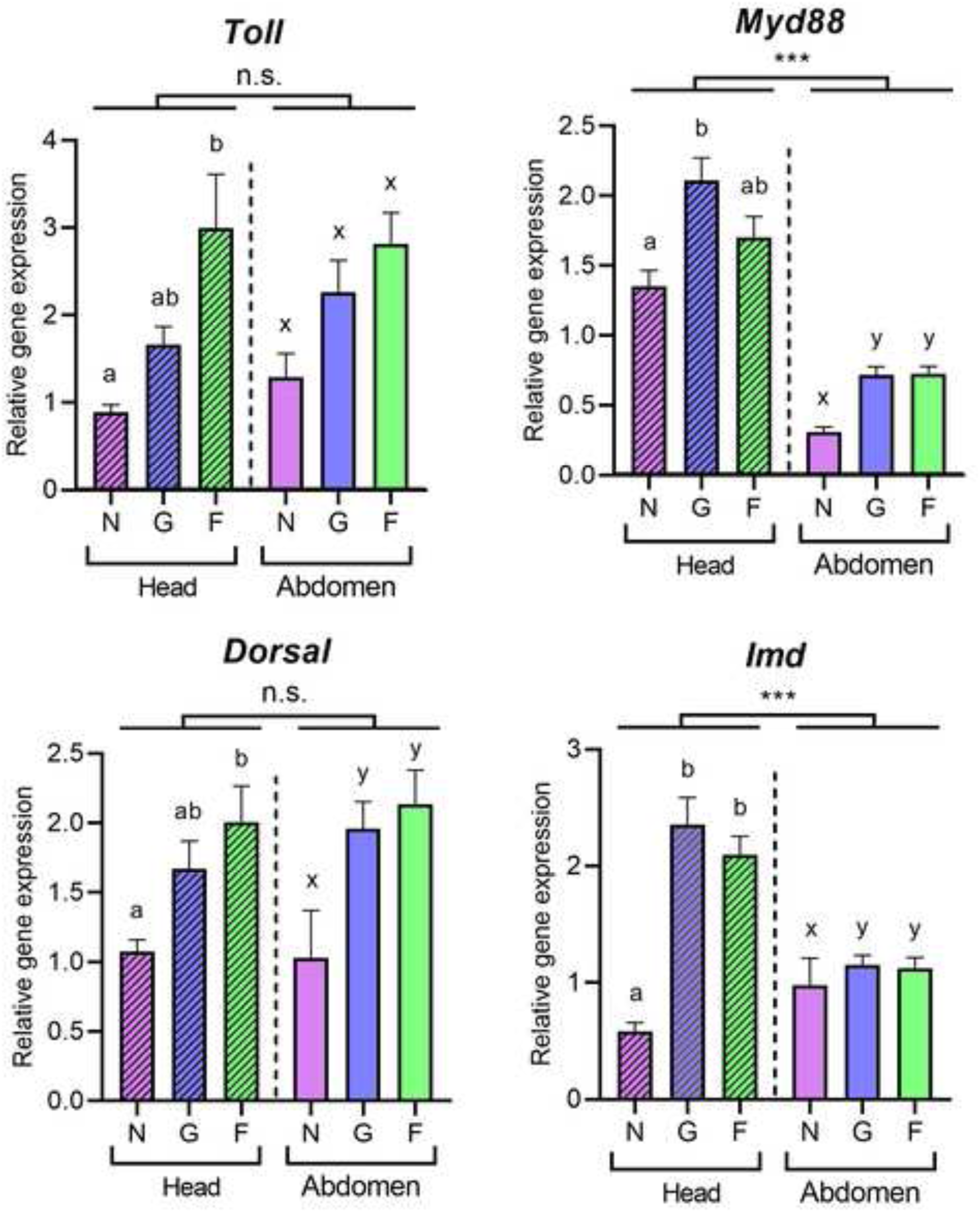
Relative expression levels of four immune system genes in the heads and abdomens of *Frieseomelitta varia* nurses, guards, and foragers. Bar plots representing the transcript levels for (A) *Toll,* (B) *Myd88*, (C) *Dorsal,* and (D) *Imd*. Purple: Nurse (N), Blue: Guard (G), Green: Forager (F). Different letters indicate statistically significantly differences among the three live cycle stages within each body compartment. Asterisks indicate statistical differences between the two body compartments (GLMM). Symbols (***) *P* < 0.0001, (**) *P* < 0.001, (*) *P* < 0.05, (n.s.) non-significant. Sample size per colony, *N* = 7; per life stage within each body compartment, *N* = 21 (3 colonies). The error bars represent the standard error of the mean (SEM).

With respect to the life cycle stages, the guards and foragers of *F. varia* and *M. quadrifasciata* in general showed a tendency towards higher levels of immunity gene expression compared to the nurse-stage workers. The workers of *M. quadrifasciata* showed differences in gene expression levels between the younger (nurse-stage) and the older bees (foragers and guards) in both body compartments for the genes *Toll* (GLMM: χ^2^ = 133.1, *P* < 0.001) and *Myd88* (Fig. 4: GLMM: χ^2^ = 4.2, *P* < 0.05). With respect to the other two genes, an abdominal up-regulation of *Dorsal* (GLMM: χ^2^ = 11.5, *P* < 0.001) was associated with the guard and forager stages, while *Imd* was higher expressed only in the heads of guards compared to nurses (GLMM: *t* = 3.2; *P* < 0.05).

For *F. varia* workers, the heads showed significantly higher *Imd* expression in both guards (GLMM: *t* = 8.25; *P* < 0.01) and foragers (GLMM: *t* = 9.75; *P* < 0.01) compared to nurses. *Dorsal* and *Toll* are both higher expressed in foragers compared to nurses (*Dorsal* GLMM: *t* = 2.8; *P* < 0.05; *Toll* GLMM: *t* = 4.57; *P* < 0.01) but not in guards, while *Myd88* is higher expressed in guards compared to nurses (GLMM: *t* = 3.12; *P* < 0.01), but not in foragers (Fig. 5; Table S5).

Notably, we found high levels of the immune gene transcript levels in the heads of guards and foragers, especially in the more aggressive species, *S. postica*. Yet, given that the abdominal fat body is the primary insect tissue for mounting an immunity response (Li et al., 2019), we asked whether these high expression levels in the bees’ heads would in fact be related to an actual immune response. To address this, we assessed the levels of *Abaecin* transcripts, as these should represent a direct readout for a canonical immune response (Lourenço et al. 2018).

The *Abaecin* gene encodes a major antimicrobial peptide, and its expression is driven by both the Toll and the Imd pathway in the honeybee (Lourenço et al., 2018). For *S. postica* and *F. varia* we found no differences in the *Abaecin* expression levels between head and abdomen (GLMM: χ^2^ = 1.8, *P* =0.17; and χ^2^ = 0.5, *P*=0.44, respectively), while *M. quadrifasciata* workers showed higher *Abaecin* expression in the abdomen than in the head (Fig. 6: GLMM: χ^2^ = 8.7, *P* < 0.003). Furthermore, guards and foragers of both *S. postica* and *M. quadrifasciata* had higher levels of *Abaecin* expression than nurses in both body compartments, while for *F. varia* and *A. mellifera,* this was the case only for the head.

**Figure 6.**
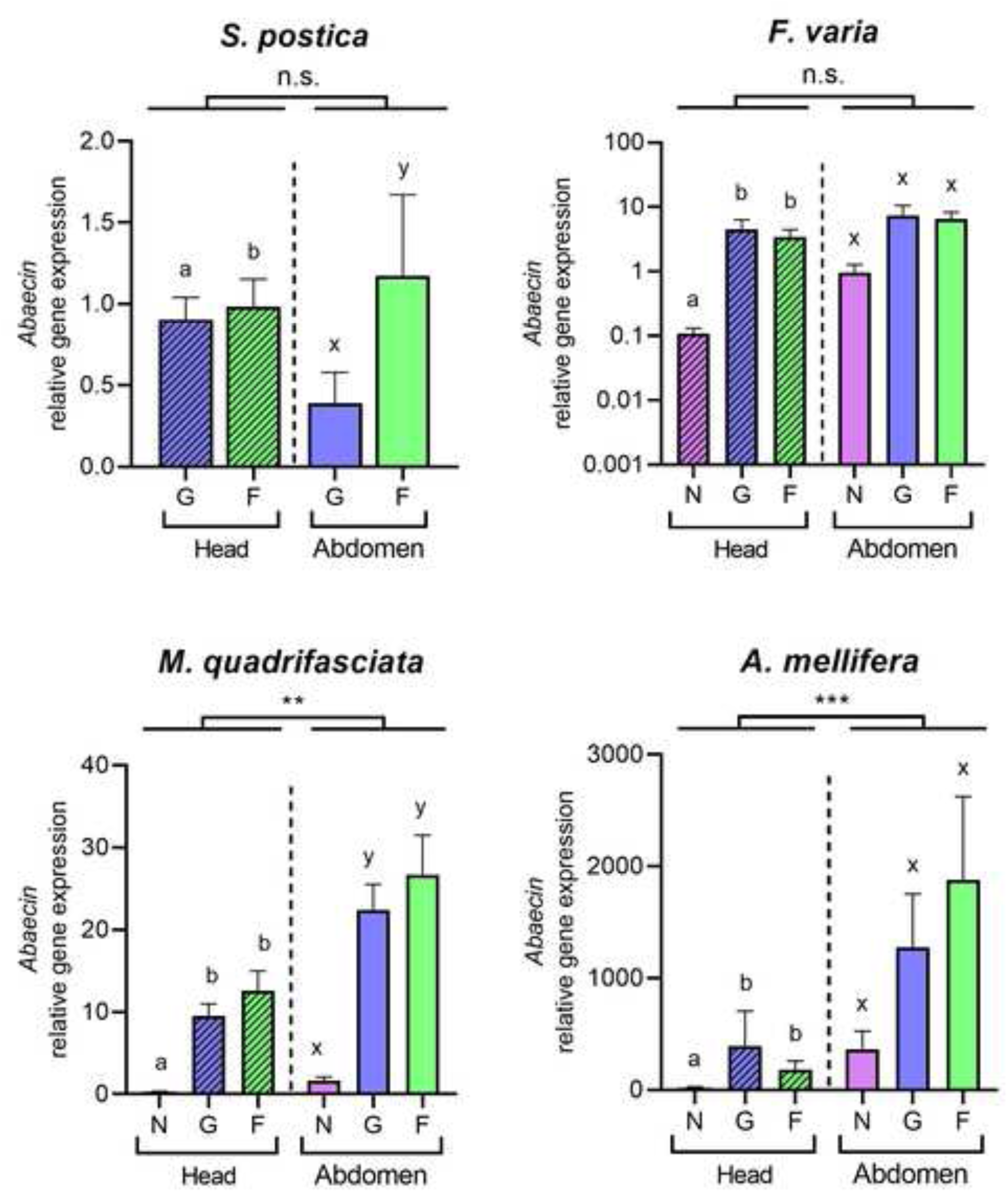
*Abaecin* gene expression in the heads and abdomens of the workers of stingless bees. Bar plots for *Abaecin* gene expression of *S. postica* (upper left) and *F. varia* (upper right) showing no differences between the two body compartments. *M. quadrifasciata* workers (lower left) and *A. mellifera* (lower right) have higher *Abaecin* gene expression levels in the abdomen. Purple: Nurse (N), Blue: Guard (G), Green: Forager (F). Different letters indicate significant differences within each body compartment. Asterisks indicate statistical differences between the two body compartments (GLMM for *S. postica*, *F. varia*, and *M. quadrifasciata*; GLM for *A. mellifera*). Symbols (***) *P* < 0.0001, (**) *P* < 0.001, (*) *P* < 0.05, (n.s.) non-significant. Sample size per colony, *N* = 7; per life stage within each body compartment, *N* = 21 (3 colonies for each stingless bee species); *N* = 7 for *A. mellifera* (one colony). The error bars represent the standard error of the mean (SEM).

The similarity in the expression levels for *Abaecin* with respect to social task status across the four species indicates that, even in the absence of an immune challenge, the immune system readiness level is higher in the older bees (guards and foragers), possibly because they perform riskier tasks outside the colony in terms of predation threat and pathogen exposure. Hence, while the expression patterns of the Toll and Imd pathway genes showed pronounced differences for the head *vs.* abdominal tissues across the four bee species, associated with their respective aggressiveness levels, the *Abaecin* expression levels showed an age/task-related pattern that was common across the four species. This discrepancy between the two signalling pathways and the actual immunity readout (*Abaecin*) indicates a non-canonical function for the Toll and Imd pathways in the heads of the more aggressive stingless bees.

### Immunity gene expression in the workers of Apis mellifera

The association between aggressiveness and immune gene expression levels in the heads of the stingless bee workers that perform the higher risk tasks of guarding and foraging, thus raised the question as to whether this could be a conserved trait across the highly eusocial corbiculate bees (Apini and Meliponini). To address this, we assessed the transcript levels of the same innate immunity genes also in the honeybee, *A. mellifera* (Fig. 7, Table S6).

**Figure 7.**
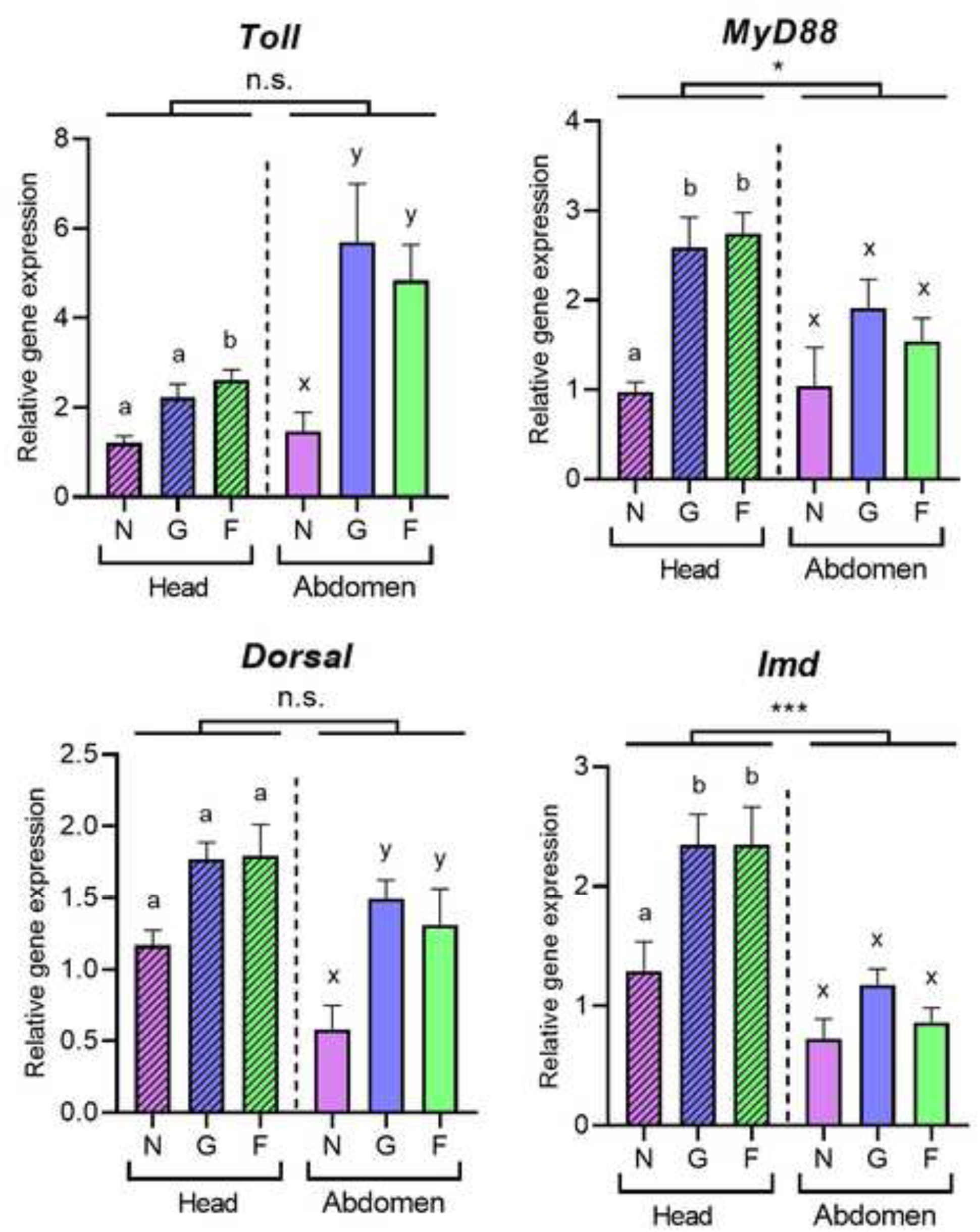
Relative expression levels of four immune system genes in the heads and abdomens of honeybee, *Apis mellifera,* workers. Bar plots representing the transcript levels for (A) *Toll,* (B) *Myd88*, (C) *Dorsal,* and (D) *Imd*. Purple: Nurse (N), Blue: Guard (G), Green: Forager (F). Different letters indicate statistically significantly differences among the three live cycle stages within each body compartment. Asterisks indicate statistical differences between the two body compartments (GLM). Symbols (***) *P* < 0.0001, (**) *P* < 0.001, (*) *P* < 0.05, (n.s.) non-significant. Sample size per life stage within each body compartment, *N* = 7. The error bars represent the standard error of the mean (SEM).

We found that only *MyD88* and *Imd* are higher expressed in the heads compared to the abdomen (GLM: χ^2^ = 4.14, *P* = 0.04; and χ^2^ = 26.86, p < 0.001, respectively). Also, for all tested genes, we observed an increase in gene expression in at least one tissue when comparing guards and foragers to nurse bees (Fig. 7; Table S6). Furthermore, like in *M. quadrifasciata*, the *Abaecin* expression was higher in the abdomen than in the head in all the life cycle stages considered in this study (GLM: χ^2^ =16.78, *P* < 0.001). These findings for the notoriously aggressive Africanized honey bee provide additional support for a possibly non-canonical function of the Toll and Imd pathways, associated with the more aggressive Meliponini species tested in this work.

## DISCUSSION

Based on the diversity in life history parameters seen in stingless bees, exemplified by the three focal species of this study, we intended to investigate the possible relationship between the bees’ social tasks, their aggressiveness, and their immune system status against the background of the worker’s reproductive biology (Fig. 1). The most striking finding was the upregulation in the expression of all four of the Toll and Imd pathway genes in the heads of the guards and foragers of *S. postica* in comparison to their abdominal tissues (Fig. 3), as this is the more aggressive among the three species that we tested (Fig. 2). In the heads of *F. varia,* only two of these genes (*Myd88* and *Imd)* are higher expressed in the heads of the workers (Fig. 5), while in *M. quadrifasciata,* the opposite is seen, with two genes (*Toll* and *Dorsa*l) actually higher expressed in the abdomen (Fig. 4). In *A. mellifera*, a possibly aggressiveness-related pattern was observed for only two of the four genes (*Myd88* and *Imd*) in the heads of guards and foragers. However, as age and task are often correlated in social insects, we acknowledge that these differences may also reflect age-related variation rather than task-specific regulation. Nonetheless, we interpret these findings as indicating a possibly non-canonical (neo)functionalization of the immune system pathways in the context of colony defence behaviours of highly eusocial bees.

### Nest defence and aggression in stingless bees

The possible reasons for the evolutionary loss of the stinging apparatus in the Meliponini are still an open question. Yet, the idea that the small size of the bees (miniaturization) would reduce the efficiency of their sting is not a very convincing argument, considering the continued presence of a sting in even smaller ants. Similarly problematic is the concomitant reduction of their venom gland, in view of the fact that genes encoding venom proteins were found conserved in stingless bee genomes (Koludarov et al., 2023). Yet, what is well known among beekeepers, is that stingless bee species greatly vary in their aggressiveness as part of their strategies of nest defence.

The results obtained here via the arena assay confirmed that *S. postica* workers, at the individual level, exhibit a strong aggression response towards an intruder. This is consistent with the fact that, among the three species that we analysed, *S. postica* also has the largest nest entrance tunnel and the largest number of guards controlling the intense flight traffic (Couvillon et al., 2008). Furthermore, when they attack people with bites, the sensation is quite painful compared to bites from other stingless bees (Shackleton et al., 2015). Hence, bees of the genus *Scaptotrigona* have an efficient, aggression-based colony defence strategy. In fact, beekeepers often emphasize the precautions that are needed when working with colonies of this genus (Blochtein, 2008).

*F. varia*, the species that showed the intermediate level of aggressiveness in the arena test (Fig. 2), also showed attention towards the intruder. However, they did not attack with bites in most of the assay rounds. They have smaller mandibles than *S. postica*, and their main defence strategy is to apply sticky resin onto intruders (Shackleton et al., 2015). This strategy may not have much of an effect against vertebrate enemies, but it is very efficient against other arthropods (Couvillon et al., 2008; Grüter, 2020). In line with that, during the arena assays the individuals were seen to perform short flights, bouncing against the intruder inside the Petri dish. In the field, such collision flights would allow the *F. varia* workers to stick the resin from their corbiculae onto the potential intruder, causing immobilization of the latter (Roubik, 2023).

In contrast to the other two species, *M. quadrifasciata* workers were completely unaggressive in the arena assay. In fact, they were even less aggressive individually than what is observed when they are confronted with a potential intruder at the entrance of a colony in the field (Breed and Page, 1991; Nunes et al., 2014).

### Pros and cons of the arena assay

The arena assay allowed us to quantitatively assess the aggressiveness level of individual stingless bee workers of three focal species, independent of variation in colony contexts. Yet, like in other behavioural tests on individual bees, certain adjustments were necessary. This included the necessity of introducing a 15-hour food restriction period prior to testing a bee’s aggressiveness towards an intruder in the arena. Like in the proboscis extension reflex (PER) assay (Bittermann et al., 1983; Smith and Burden, 2014) a food restriction turned out to be necessary for the appropriate motivation of the bees to respond to the stimulus. As previously shown in a different behavioural context, food restriction affects the metabolism of *A. mellifera* workers, including their brain metabolism (Rittschof et al., 2018). Furthermore, there is strong evidence that aggressive behaviours are associated with significant metabolic changes in the brains of honeybees (Chandrasekaran et al., 2015; Li-Byarlay et al., 2014), underlining the importance of taking into account the bees’ nutritional status when designing and standardising aggression tests.

With this in mind, there is the possibility that bees of larger size, like *M. quadrifasciata,* may have larger energy reserves than the smaller *S. postica* and *F. varia*, and potentially, this could make them less responsive in the arena assay. However, when we extended the food restriction to 24 hours, this led to increased mortality. Hence, the results of the arena assay performed under these standardised conditions confirmed the field observations regarding the differences in the species’ aggressiveness also at the level of the individual workers.

Notably, we did not perform the arena test for the testing of honeybee workers, because the protocol conditions that we used for the stingless bees were found not suitable for the former. The main reason was that for the confinement in the cages, the stingless bee workers received a diet that contained pollen. Yet, *A. mellifera* workers consuming pollen hardly survive for more than 14 days in the cages because, different from the stingless bees, they cannot defecate in confinement, without flying (Cole et al., 2021). Furthermore, an arena aggression assay for *A. mellifera* workers was formerly designed and performed with small groups of bees and not with individual bees (Nouvian et al., 2016).

### Elevated gene expression levels in the head of an aggressive stingless bee species indicate a non-canonical function for innate immunity signalling pathways

To address the question as to whether and how the species-specific levels of aggressiveness and the innate immune system of the stingless bee workers may be intertwined in terms of life history strategies and trade-offs (i.e., reproduction vs. immunity/somatic maintenance), we sampled workers according to their social task in the colony. What we found is that in the more aggressive species, *S. postica*, the levels of immune gene expression were upregulated in the head of the guards and foragers (Fig. 3). Since the abdominal fat body, but not the head, is the primary organ responsible for the production of antimicrobial peptides (Li et al., 2019), this strong overexpression of the Toll and Imd pathway genes in the bees’ heads is clearly remarkable.

*Toll* and *Dorsal* genes (*NF-kB* in vertebrates) are present in the genomes of most organisms, and the Toll/Toll-like receptors (TLR) signalling pathway is a very ancient component in metazoan history (Leulier and Lemaitre, 2008). Actually, while it has been hypothesized that the TLR immune functions involving Myd88 and NF-kB arose in the Bilateria (Kim and Ausubel, 2005), all the core elements of Toll signalling are already present in Cnidaria (Chapman et al., 2010). And even though the primary function of Toll signalling is the recognition of pathogenic microorganisms, it has also acquired an embryonic function in insects, determining the dorsoventral axis (Anderson et al., 1985). Furthermore, in addition to its canonical function as a transmembrane receptor for immune signalling, Toll may serve additional roles in the nervous system in both vertebrates and invertebrates (Dresselhaus and Meffert, 2019).

Myd88, an adapter protein downstream to the transmembrane protein Toll (Horng and Medzhitov, 2001), exhibited higher expression levels in the heads of *S. postica* guards compared to foragers (Fig. 3). Interestingly, Myd88 is already known to play an important regulatory role in mammalian neural and behavioural functions, including the regulation of chronic stress responses (Hosoi et al., 2021). Furthermore, inhibition of this protein promotes depression-like behaviour (Yao et al., 2023) and depression-like sleep disorders (Choudhury et al., 2022). Since *Myd88* was found higher expressed in the aggressive guard bees of *S. postica,* this molecular signature strengthens the hypotheses that ancestral immunity proteins were co-opted to shape complex collective colony defence behaviours in the highly eusocial stingless bees.

Dorsal (NF-kB), the main transcription factor of the Toll pathway, was also found overexpressed in the heads of *S. postica* workers compared to their abdomens, whereas in the docile *M. quadrifasciata* workers, this pattern is inverted. In mammals, NF-kB is known to be involved in cognitive processes (Dresselhaus and Meffert, 2019). It is constitutively activated in neurons from various regions of the central nervous system, mainly in glutamatergic neurons (Bhakar et al., 2002; Kaltschmidt et al., 1994), and it notably accumulates in the terminal regions of these neurons. Compelling evidence indicates that the activation of NF-kB is induced by the neurotransmitter glutamate and the influx of calcium ions (Ca^2+^) (Lilienbaum and Israël, 2003; Meffert et al., 2003), and upon activation, NF-kB undergoes a retrograde transport and becomes translocated to the nucleus of the respective neurons (Meffert et al., 2003; Meffert and Baltimore, 2005; Wellmann et al., 2001). In this way, it functions as a transducer of synaptic information to the nucleus, where it assumes its role as a transcription factor (Mémet, 2006). In addition, NF-κB is associated with memory consolidation and learning in vertebrates and invertebrates, such as crustaceans and insects (Freudenthal and Romano, 2000; Mémet, 2006; Merlo et al., 2005). This response apparently occurs in neurons, but not in glial cells, where NF-κB is strictly associated with the control of mammalian immune responses and metabolism (Dresselhaus and Meffert, 2019). Based on our results obtained for stingless bees, it is thus, reasonable to assume that in these bees, the Toll signalling pathway, including its key transcription factor Dorsal, were co-opted to perform a behavioural function in addition to its canonical immune system role.

With respect to *Imd* expression, the pattern was similar to *Dorsal* in *S. postica* in terms of body compartment and social task specificity. However, aside from its immune system function, little is known about potentially alternative roles of Imd signalling in insects. Nonetheless, there are reports on its involvement in neurodegeneration, cell death, and regulation of the gut microbiota (Hanson and Lemaitre, 2020; Zhai et al., 2018). In this respect, our results may be the first ones to identify a link between the Imd pathway and behavioural traits also in an invertebrate.

As emphasized, the expression patterns of all four immune signalling pathway genes quantified in this study indicate an increase associated with the age-related social task performed by the workers of the stingless bees, in both the head and the abdomen. But how does this actually relate to the bees’ immune system status? In *D.* melanogaster, healthy and free of infection individuals showed a chronic increase with age in the immune system activity, leading to an increased expression of antimicrobial peptides (AMPs) in the brain, followed by marked neurodegeneration (Stuart et al., 2022). Furthermore, a study by Kounatidis et al. (2017) showed that suppression of the Imd/NF-κB signalling pathway in glial cells resulted in an extension of the flies’ lifespan. This is in line with data for *A. mellifera* (Lourenço et al., 2019), where nurse bees have much lower expression levels of immunity genes and an increased lifespan.

In adult insects, AMPs are the most prominent products generated by the activation of the Imd and Toll signalling pathways (Bulet and Stocklin, 2005), and activation of the immune system in the brain leads to high AMP expression. This can have collateral cytotoxic effects related to neurodegeneration, and consequently behavioural changes, including sleep and memory dysregulation (Barajas-Azpeleta et al., 2018; Stączek et al., 2023). The brain of *A. mellifera* was previously seen to exhibit AMP overexpression upon infection with Deformed Wing Virus, a common bee virus, demonstrating the ability of the brain to elicit a strong immune response (Pizzorno et al., 2021). Studies with *D. melanogaster* in turn demonstrated that the production of AMPs may also be regulated by pathways other than Toll and Imd (Stączek et al., 2023). For instance, the transcription factor FoxO, which is a downstream regulator of the insulin signalling pathway, can activate the expression of AMPs in flies undergoing starvation, even in the absence of an infection or injury event (Becker et al., 2010). Therefore, the elevated *Abaecin* transcript levels that we found in the head of the older stingless bees may not be exclusively driven by the immune system pathways Toll and Imd, but also by pathways associated with their age-specific diets and, consequently, metabolic regulation. Nonetheless, in our previous study, where we compared head and abdominal expression of candidate genes for social task performance in stingless bees (de Souza and Hartfelder, 2023) we did not see any expression difference in the abdominal *tor* gene expression for *F. varia* and *M. quadrifasciata* nurses and foragers. In their heads, however, we saw an inverse expression pattern between the two species, indicating that this major nutrient-sensing pathway may be linked to the head immune gene expression profiles.

Finally, in this respect as already previously stated, we recognize as a limit of our study that we cannot not separate between age and social function. These two factors are strongly intertwined, with young bees preferentially performing tasks within the nest, while older ones take on guarding and foraging task, and these tasks are strongly associated with immune system functionality (Amdam et al., 2005; Lourenço et al., 2019).

### Immune system gene expression and the life histories of social bees

At this point it is important to state that the question that we asked differs from the one addressed in most studies on immune system functions in social insects. Specifically, we did not ask how does the bees’ immune system respond to a pathogen challenge, but rather, whether the aggressiveness level of the workers, as part of the colony defence against intruders and predators, is reflected in their (unchallenged) immune system status. We addressed this question in the workers of three species of stingless bees that vary in important life history traits, not only with respect to nest defence, but also with the workers’ reproductive status.

Different from the honeybees, whose workers are functionally sterile in the presence of the queen, the workers of stingless bees differ considerably in their potential of laying eggs (Hartfelder et al., 2006; Vollet-Neto et al., 2018). While *F. varia* workers are completely sterile, those of *M. quadrifasciata* and *S. postica* lay trophic and also reproductive eggs in the presence of a queen, thus contributing to the colonies’ reproductive output. A key reproductive protein in female insects is vitellogenin (Vg) (Sappington and Raikhel, 1998), but interestingly, in the functionally sterile honeybee workers, the major function of Vg is not oogenesis. Rather, together with juvenile hormone, it regulates the behavioural transition from intranidal (nursing) to extranidal (foraging) behaviour (Amdam and Omholt, 2003), and it is in this context that the high Vg levels in the haemolymph of the nurses prevent the onset of immunosenescence (Amdam et al., 2005). Moreover, Vg itself is a powerful antioxidant protein that plays key roles in the regulation of the lifespan of the honeybee queens and workers (Corona et al., 2007; Havukainen et al., 2013; Seehuus et al., 2006).

In a recent study on gene expression underlying the nurse-to-forager transition in stingless bees (de Souza and Hartfelder, 2023) we found an exactly opposite expression pattern for the *Vg* gene in two species, showing upregulated transcript levels in the heads of *F. varia* foragers, which are sterile, contrasting with upregulated *Vg* levels in the abdomen of *M. quadrifasciata* nurses, which can lay eggs. Now taken together with the current data on immunity gene expression, this suggests that, contingent on the reproductive status of the workers, there appears to be a trade-off in the investment between the Vg-related and the Toll/Imd-related immunity status. This trade-off not only extends beyond the honeybees (Lourenço et al., 2012), it also appears to be species-specifically balanced against the aggressiveness level in the stingless bees, as shown in the current study.

## Conclusion

The high transcript levels of the Toll and Imd pathway genes in the heads of the workers of the aggressive species *S. postica*, compared to those in the docile workers of *M. quadrifasciata*, indicate a possible non-canonical involvement of the innate immune system in behaviours related to nest-defence strategies of social bees. The nest is an important social factor that protects the society and its individuals from stressful situations and, inherently, shapes their life histories (Walton et al., 2014). Given the intricate interplay between the innate immune system and stress responses documented in insects (Štětina et al., 2019; Hao et al., 2023), including bees (McMenamin et al., 2020), made us, thus, raise the hypothesis of a possible association between the patterns of immune gene expression and the levels aggressiveness in the workers of social bees.

Of specific interest in this context is the recent proteomics study by McAfee et al. (2024), who investigated a possibly caste-specific reproduction-immunity trade-off in the bumble bee *Bombus impatiens*. While they found no evidence for such a trade-off in the workers, they did so for the queens. However, the authors consider this to be due to confounding effects resulting from the queens’ complex life cycle, including a diapause phase for overwintering, followed by a solitary nest foundation before a functional colony gets established. They conclude that additional factors, such as worker age and aggressiveness should be taken into consideration. It is this question what we now addressed in the stingless bees. Not only do their queens spend their entire life in the centre of a well-protected nest, except for a brief mating flight, it is also the species richness of the Meliponini and their extraordinary variability in the biology of their workers that makes them a valuable source for comparative studies on life history strategies and functional genomics in social bees.

As next a step, extending this study to other stingless bee species that aggressively defend their nests, or aggressively perform raids on other colonies, like *Lestrimelitta limao*, will be important to understand whether this non-canonical pattern of immune system gene expression represents a general trait in these highly eusocial bees. Furthermore, the development of behavioural assays that allow the quantification of aggressiveness levels, such as the arena assay, coupled with pharmacological manipulation and/or RNAi functional assays, should allow mechanistic insights into the importance of the Toll and Imd pathways in the sophisticated social behaviours of the bees.

## DATA AVAILABILITY

All relevant data can be found within the article and its supplementary information. The raw data is available from the following link: https://doi.org/10.5281/zenodo.13374927

## ABBREVIATIONS

AMP: Antimicrobial peptide
GLM: General Linear Model
GLMM: General Linear Mixed Model
NF-kB: Nuclear Factor-kappa B
RT-qPCR: Quantitative reverse transcription polymerase chain reaction
TLRs: Toll-like receptors
Vg: Vitellogenin

## ACKNOWLEDGMENTS

We thank Jairo de Souza and Luiz Roberto Aguiar for helping with the maintenance of the bee colonies in the meliponary and apiary of the Ribeirão Preto Campus of the University of São Paulo.

## SUPPLEMENTARY MATERIAL

**Figure S1** – Effect of dietary restriction on the number of attacking and non-attacking workers.

**Figure S2** – Images illustrating the phenotypic characteristics of the workers of the three stingless bee species used in this study

**Figure S3** – Ct values of the two reference genes (*Rps18* and *Rpl32*) across behavioral subcastes and tissues in the *S. postica*, *F. varia* e *M. quadrifasciata*.

**Table S1** - Primers used in the RT-qPCR assays.

**Table S2** - Indices of aggressiveness for individual workers of stingless bees tested in arena assays.

**Table S3** - Statistical details of the GLMM test used to analyse the expression of immune pathways genes in *S. postica*.

**Table S4** - Statistical details of the GLMM test used to analyse the expression of immune pathways genes in *M. quadrifasciata*.

**Table S5** - Statistical details of the GLMM test used to analyse the expression of immune pathways genes in *F. varia*.

**Table S6** - Statistical details of the GLM test used to analyse the expression of immune pathways genes in *A. mellifera*.

